# Joint models reveal human subcortical underpinnings of choice and learning behaviour

**DOI:** 10.1101/2024.12.20.629647

**Authors:** Steven Miletić, Niek Stevenson, Pierre-Louis Bazin, Anneke Alkemade, Scott J. S. Isherwood, Anne C. Trutti, Desmond H. Y. Tse, Asta K. Håberg, Birte U. Forstmann

**Author notes:** These authors contributed equally to this work (Steven Miletić, Niek Stevenson).

## Abstract

Decision making and learning processes together enable adaptive goal-oriented behaviour. Animal studies demonstrated the importance of subcortical regions in these cognitive processes, but the human subcortical contributions remain poorly characterised. Here, we study choice and learning processes in the human subcortex, using a tailored ultra-high field 7 T fMRI imaging protocol combined with joint models. Joint models provide unbiased estimates of brain-behaviour relations by simultaneously including behavioural and neural data at the participant and group level. Results demonstrate relations between subcortical regions and the adjustment of decision urgency. Value-related BOLD differences were found with opposite BOLD polarity in different parts of the striatum. Multiple sub-cortical regions showed BOLD signatures of reward prediction error processing, but contrary to expectations, these did not include the dopaminergic midbrain. Combined, this study characterises the human subcortical contributions to choice and learning, and demonstrates the feasibility and value of joint modelling in facilitating our understanding of brain-behaviour relationships.

## 1 Introduction

Decision making and instrumental learning continuously interact (Sutton & Barto, 2018): Error-driven learning processes refine and update the information on which value-based choices are made. In behavioural studies, recent advances have integrated insights from the traditionally separate fields of perceptual decision-making on the one hand, and error-driven learning on the other, into a singular framework (Fontanesi, Gluth, Spektor, & Rieskamp, 2019; Fontanesi, Palminteri, & Lebreton, 2019; McDougle & Collins, 2020; Miletić, Boag, & Forstmann, 2020; Miletić et al., 2021; Pedersen & Frank, 2020; Pedersen, Frank, & Biele, 2017; Sewell, Jach, Boag, & Van Heer, 2019; Sewell & Stallman, 2020; Shahar et al., 2019; Turner, 2019; Wagner, Mathar, & Peters, 2022). The combination of evidence accumulation to threshold (a core principle from decision-making research) and simple delta rules (a core principle in reinforcement learning) was shown to provide a precise characterisation of behaviour in instrumental learning tasks: It can explain response time distributions, choice accuracy, and the learning-related changes in response time distributions and choice accuracy.

While providing a rich account of the algorithmic processes underlying choice and learning, cognitive models are agnostic about the neural implementation, which is our focus here. Both fields can lean on rich literatures on the relation between neural and behavioural data, although based largely on animal recordings. In decision making, the basal ganglia have long been implicated in action selection (Chevalier, Vacher, Deniau, & Desban, 1985; Deniau & Chevalier, 1985; Frank, Loughry, & O’Reilly, 2001; Lo & Wang, 2006; Mink, 1996; Mink & Thach, 1993; Nambu et al., 2000; Redgrave, Prescott, & Gurney, 1999). Furthermore, key insights were obtained from recordings that demonstrated processes resembling evidence accumulation in a variety of brain regions including the basal ganglia (Ding & Gold, 2010; Lauwereyns, Watanabe, Coe, & Hikosaka, 2002; Thura & Cisek, 2017), the superior colliculus (Basso & Wurtz, 1998; Grimaldi, Cho, Lau, & Basso, 2018; Horwitz & Newsome, 1999; Jun et al., 2021; Munoz & Wurtz, 1995; Ratcliff, Cherian, & Segraves, 2003; Ratcliff, Hasegawa, Hasegawa, Smith, & Segraves, 2007), and cortical regions including parietal cortex (Churchland, Kiani, & Shadlen, 2008; Huk & Shadlen, 2005; Kiani, Hanks, & Shadlen, 2008; Mazurek, Roitman, Ditterich, & Shadlen, 2003; Roitman & Shadlen, 2002; Shadlen & Newsome, 1996, 2001; Steinemann et al., 2024), the frontal eye fields (Ding & Gold, 2012; Hanes & Schall, 1996; Kim & Shadlen, 1999; Mante, Sussillo, Shenoy, & Newsome, 2013; Purcell et al., 2010; Schall, 2002; Thompson, Hanes, Bichot, & Schall, 1996), and premotor and motor cortex (Cisek & Kalaska, 2005; Peixoto et al., 2021; Romo, Herńandez, & Zainos, 2004; Thura & Cisek, 2016). In parallel, studies in reinforcement learning have long focused on the role of the dopaminergic midbrain in calculating reward prediction errors, and on dopamine as a signal conveying reward prediction errors (e.g., Gershman et al., 2024; Montague, Dayan, & Sejnowski, 1996; Schultz, 1986; Schultz, Apicella, & Ljungberg, 1993; Schultz, Dayan, & Montague, 1997; Watabe-Uchida, Eshel, & Uchida, 2017).

Thus, both fields suggest prominent involvement of subcortical regions. Unfortunately, in humans, the role of subcortical regions in decision making and learning is less well characterised. This is due to various factors that make imaging the sub-cortex particularly difficult. Many subcortical regions suffer from signal losses when conventional functional magnetic resonance imaging (fMRI) methods are used. The underlying causes include the deep location of the subcortex, high iron concentrations, and the small sizes of individual regions (for an overview, see Forstmann, De Hollander, Van Maanen, Alkemade, & Keuken, 2017). Because of these factors, the majority of human neuroimaging studies have focused on the neocortical sheet, combined with the larger subcortical regions including the striatum and thalamus (for a meta-analysis, see Keuken, Van Maanen, Boswijk, Forstmann, & Steyvers, 2018). To achieve the signal quality necessary for investigating the typically small blood-oxygenation level dependent (BOLD) responses associated with cognitive functions in smaller regions, specialised MRI protocols designed at ultra-high field strengths of 7 Tesla (T) have been developed (De Hollander, Keuken, van der Zwaag, Forstmann, & Trampel, 2017; Miletić, Bazin, et al., 2020; Miletić, Keuken, et al., 2022).

Signal quality is not the only factor to consider when discussing the challenges of studying the human subcortex. Statistical considerations form a second factor hampering the characterisation of the human subcortex in cognitive processes. Model-based analysis methods offer a principled advantage in terms of bridging the algorithmic and neural levels of analysis (Teller, 1984; Turner, Palestro, Miletić, & Forstmann, 2019). Traditional model-based MRI studies, however, rely on *two-stage* approaches, in which a cognitive model is first fit to behavioural data, and the resulting parameters are used as regressors in the analysis of the neural data. While straightforward to implement, two-stage approaches do not fully take into account the reciprocity in the relation between behaviour and the brain. Using this approach, the neural model is informed by the cognitive data, but the cognitive model is not informed by the neural data. Furthermore, the measurement uncertainty in the parameters of the cognitive model are ignored. When unaccounted for, this source of noise causes negatively biased effect sizes, a phenomenon known as attenuation (Gelman & Hill, 2006; Spear-man, 1904). It also comes at the risk of overconfidence in the effects of covariates, since the uncertainty in the estimation of the covariate is ignored (Gelman & Hill, 2006). This is especially detrimental when studying noisy data such as fMRI time-series obtained from the human subcortex. Joint models, which simultaneously model both the neural and behavioural modalities of data, at all levels of the hierarchy (participant and group level), are required to remedy this issue and achieve full statistical power (Turner, Forstmann, Love, Palmeri, & Van Maanen, 2017; Turner, Forstmann, & Steyvers, 2019; Turner, Palestro, et al., 2019; Turner, Rodriguez, Norcia, McClure, & Steyvers, 2016; Turner, Wang, & Merkle, 2017).

This study takes a joint modelling approach to studying decision-making processes and instrumental learning in the human subcortex. We bring together three contributions. Firstly, we use a single task paradigm combined with a single cognitive model, that unifies the study of decision making and reinforcement learning processes (Miletić, Boag, & Forstmann, 2020; Miletić et al., 2021), and allows for disentangling potential interactions between decision making and learning. In this task, participants are required to repeatedly make value-based choices between abstract symbols, and learn from the probabilistic reward associated with each symbol. Prior to each choice, participants are informed to emphasise either response speed or choice accuracy, thereby enforcing a change in choice strategy.

Secondly, we used an fMRI protocol tailored to meet the specific requirements for studying small subcortical nuclei at an ultra-high field of 7 T (De Hollander et al., 2017; Groot et al., 2023, 2024; Isherwood et al., 2023; Lloyd et al., 2024; Miletić, Bazin, et al., 2020; Miletić, Keuken, et al., 2022; Trutti et al., 2024). This protocol includes a short echo time to match the low T2* of iron-rich nuclei, small voxels to mitigate partial voluming effects, and a relatively high repetition time. Furthermore, we acquired multimodal quantitative anatomical MRI data, which enabled us to delineate individual subcortical nuclei with automated algorithms (Bazin, Alkemade, Mulder, Henry, & Forstmann, 2020).

Finally, we analysed brain-behaviour relations in the resulting data using high-powered Bayesian joint modelling techniques, in which two reciprocal links between neural and behavioural data are included: Reward prediction errors and value estimates of the reinforcement learning model are fed forward to the neural models within subjects, and simultaneously, across participants, inter-individual correlations between neural and behavioural model parameters are estimated. The simultaneous estimation of the cognitive and neural models allows for all sources of uncertainty to be modelled accurately, which leads to unbiased estimates of the brain-behaviour relations.

## 2 Results

Thirty-seven participants performed an instrumental learning choice task (Figure 1A) while undergoing 7 T BOLD-fMRI. They made repeated decisions between two abstract choice symbols, each followed by choice-dependent probabilistic rewards, which they used to inform subsequent choices. In total, each participant made 342 decisions. Prior to each decision, participants were instructed to emphasise either response speed or response accuracy. The behavioural data, consisting of response times and choices, were modelled with the reinforcement learning-advantage racing diffusion (RL-ARD) model (Miletić et al., 2021). This model proposes that decisions are formed through an evidence-accumulation process, where the rate of accumulation depends on the sum of an urgency signal and the internal representations of the value of each choice option (Figure 2A). The values of choice options are learned via a standard delta rule (Rescorla & Wagner, 1972). The effect of the speed and accuracy instructions were modelled by allowing both the urgency and threshold parameters to vary with instructions, in line with previous work (Miletić et al., 2021). Threshold refers to the overall amount of evidence that participants require to inform their decisions, whereas urgency refers to how participants become less patient as within a trial as time passes. The RL-ARD provided an adequate account of the behavioural data, capturing the learning-dependent increase in accuracy, decrease in response time, and the differences in RT and choice accuracy between the speed-emphasised and accuracy-emphasised trials (Figure 1B).

**Fig. 1.**
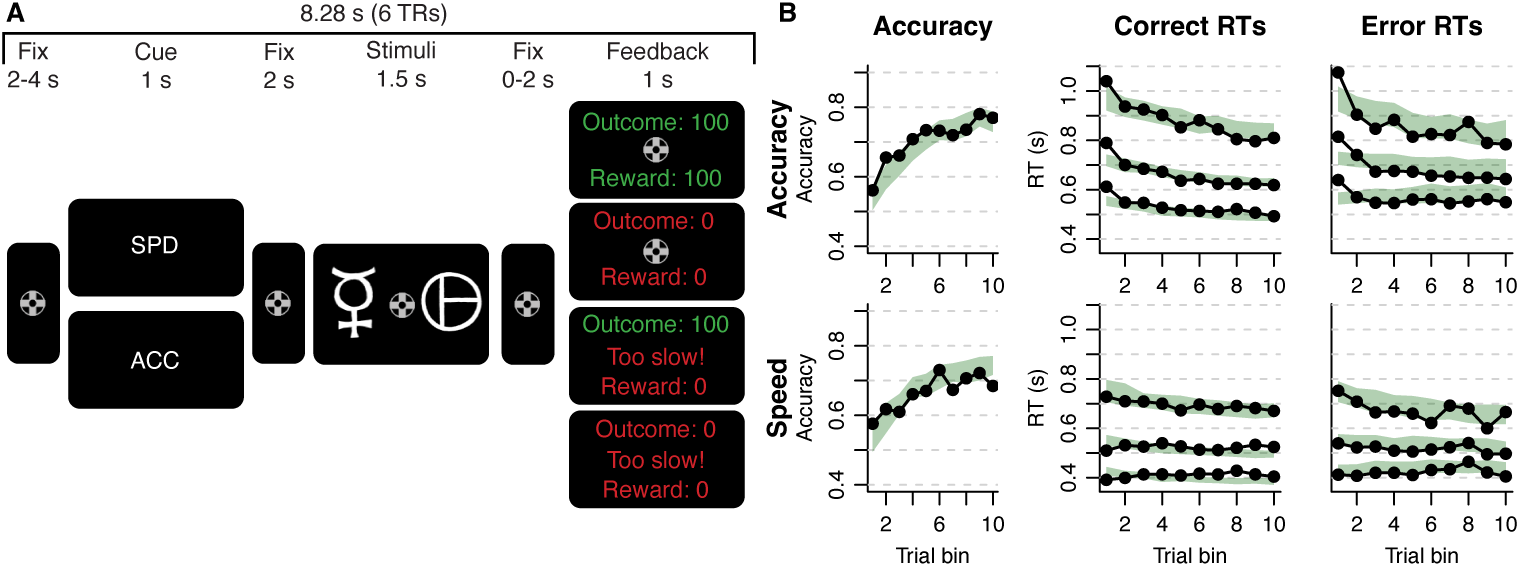
A. Experimental paradigm. Each trial started with a fixation cross, followed by the speed– accuracy trade-off cue (‘SPD’ or ‘ACC’), another fixation cross, the stimuli representing choice options, another fixation cross, and feedback. Feedback depended both on the response time (in time or too slow) and on the outcome of the probabilistic gamble (0 or 100 points). Rewards were only given if the response was in time. Durations of the fixation crosses were jittered to decorrelate event timing. B. Data (black) and model fit (green) of the RL-ARD model in the accuracy (top) and speed (bottom) condition. Left column depicts accuracy over trials, where trials were binned into 10 bins. Middle and right panel show 10th, 50th, and 90th RT percentiles for the correct (middle) and error (right) response over trial bins. Shaded areas in the middle and right column correspond to the 95% credible interval of the model fit.

**Fig. 2.**
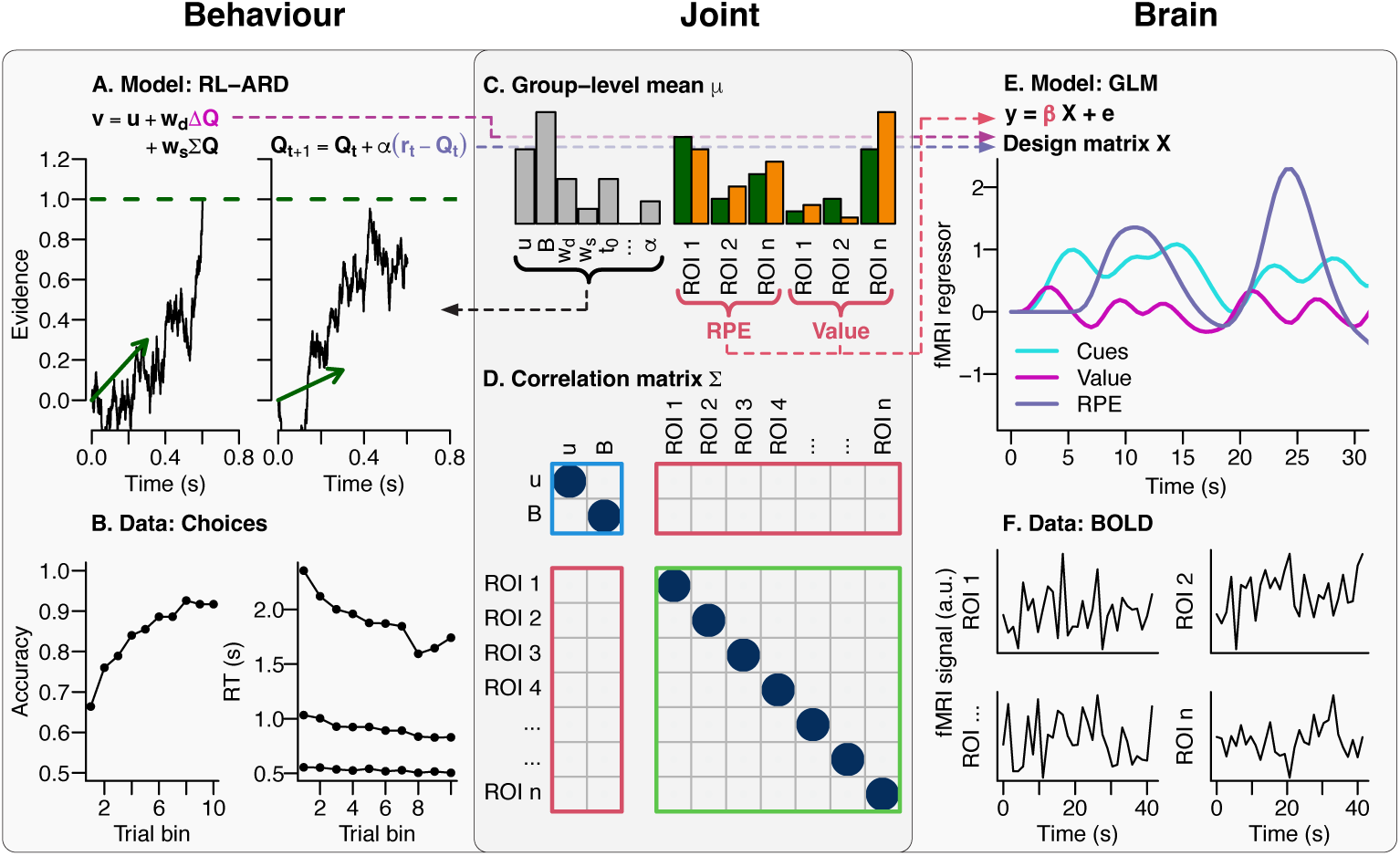
Overview of joint modelling approach. The behavioural model (A) is informed by the RT and choice data (B; see Figure 1B for detail on the visualisation of the behavioural data). The trial-by-trial differences in Q-values and prediction errors are fed forward to the design matrix of the GLM (E). The GLM is informed by the neural data (F). Mutual constraint between the two modalities of data is enabled by the joint structure that uses a multivariate normal distribution at the group level. This is described by a group-level mean (C) and correlation matrix (D). All behavioural and neural parameters are estimated simultaneously on the group level and participant level. Brain-behaviour associations of reward prediction errors and value differences are charac-terised with group-level means, while brain-behaviour associations between speed-accuracy trade-off behaviour and neural responses are estimated as inter-individual correlations. The correlation matrix is divided into behaviour-behaviour correlations (blue rectangle), brain-brain correlations (green), and brain-behaviour correlations (red). For visualisation purposes, only a subset of the parameters are shown.

In a separate session, participants underwent high-resolution quantitative MRI scans that allowed us to derive multimodal anatomical data (T1 maps, T2* maps, and quantitative susceptibility maps), which were used to delineate 17 subcortical regions of interest using the multi-contrast anatomical subcortical structure parcellation (MASSP) algorithm at the individual level (Bazin et al., 2020). The masks of the gray matter structures — the amygdala (Amg), claustrum (Cl), globus pallidus interna (GPi) and externa (GPe), periaqueductal gray (PAG), pedunculopontine nucleus (PPN), red nucleus (RN) substantia nigra (SN), subthalamic nucleus (STN), striatum (Str), thalamus (Tha), and ventral tegmental area (VTA) — were subsequently used to extract timecourses of the signal from the fMRI data. Figure 3A provides an overview of these ROIs.

**Fig. 3.**
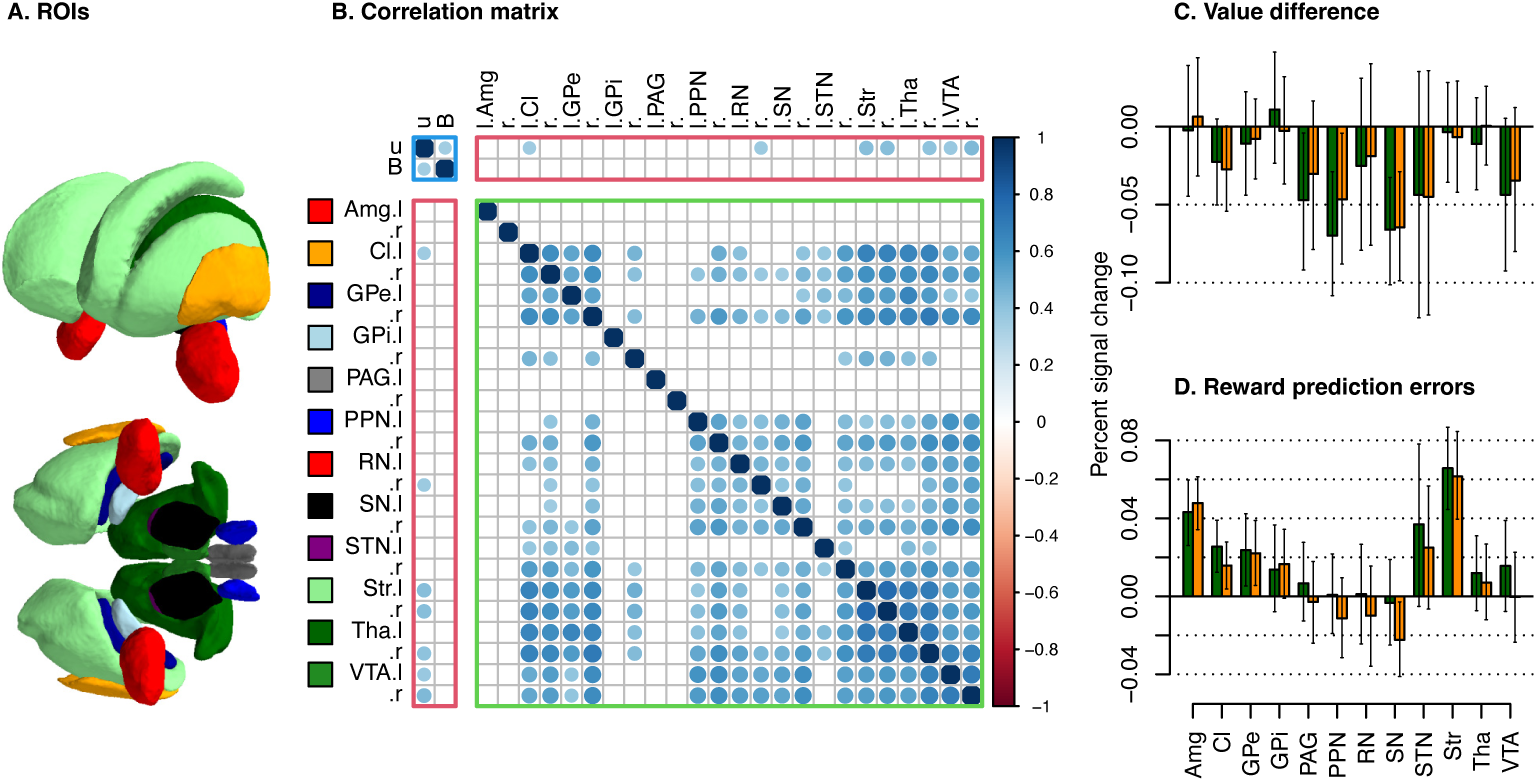
Joint model fit to the MASSP ROIs. A. Illustration of the ROIs, viewed from the front-left (top) and bottom (bottom). B. Group-level correlation matrix, which is split into behaviour-behaviour relations (outlined by a blue rectangle), brain-brain relations (red), and brain-behaviour relations (green). Only credible correlations are shown; non-credible correlations are displayed as empty squares. Relations are considered credible when the 95% credible interval of the correlation coefficient does not cover 0. All parameters are related to the speed-accuracy trade-off contrast: Its effect on urgency (u), threshold (B), and the BOLD contrast in the ROIs. C and D. Group-level estimates of within-participant brain-behaviour relations of value learning and reward prediction errors. Barplots show the percentage signal change per unit change in value difference (C) and reward prediction errors (D), for each region of interest. Green and orange bars depict the left and right hemisphere, respectively. Error bars indicate 95% credible intervals.

These neural fMRI timecourses were modelled with a general linear model (GLM; Figure 2E) which, next to a set of nuisance regressors (see Methods), included cues (speed and accuracy), stimulus value differences, and reward prediction errors, as regressors of interest. The latter two regressors were derived from the RL-ARD model, and vary across trials within participants. We estimated their mean effect on the group level (Figure 2C). We also estimated the correlations between the speed–accuracy con-trasts in the neural models (one per region of interest) and speed–accuracy difference between the urgency and threshold differences as derived from the RL-ARD (Figure 2D). Combined, this resulted in three brain-behaviour relations per region of interest that were jointly informed and reciprocally constrained by the two modalities of data.

The resulting joint model is visualised in Figure 3. Figure 3B shows the inter-individual correlations between strategic adjustments in choice behaviour (urgency and threshold) and the BOLD responses in the subcortical regions (see Table A1). The joint models revealed correlations between urgency and neural responses bilaterally in the Str and VTA, left Cl, and right RN and Tha. Next, we turned to brain-behaviour relations of value learning. The PPN and SN showed relations with value differences, as well as the left PAG (Figure 3C). The joint model further indicated reward prediction error processing in the Amg, Cl, GPe, and Str (Figure 3D). Interestingly, we found no evidence for involvement of the VTA or SN in reward prediction error coding; if anything, results indicated a *negative* association between reward prediction errors and neural activity in the right SN.

The results so far indicated involvement of the Tha (as a single region covering all nuclei) in the speed-accuracy trade-off. In a second joint model, we zoomed in on the individual thalamic nuclei using a thalamus atlas (Iglesias et al., 2018). Here, we focused only on regions larger than 150 mm^3^ in both hemispheres: the anteroventral (AV), centromedian (CM), lateral posterior (LP), mediodorsal (MD), pulvinar, ventral anterior (VA), ventral lateral (VL), and the ventral posterolateral (VPL) nucleus. In the atlas, the MD is split into a lateral and medial part (MDl, MDm), the pulvinar in an anterior, inferior, lateral, and medial part (PuA, PuI, PuL, PuM), and the VL in an anterior and posterior part (VLa, VLp). Figure 4A illustrates the ROIs that were included. The joint model based on thalamic nuclei highlighted that the brain-behaviour correlations with speed-accuracy trade-off settings were found bilaterally within the AV, CM, MDm, PuA and PuM, as well as in the right LP, VLa, and VLp (Figure 4B, see Table A2). Again, these correlations are with urgency, and appear to dominate in the right hemisphere. In the thalamic regions, we found evidence for a relation with value difference only in the right VPL (Figure 4C). Evidence for reward prediction error processing was found in the CM, PuI, and VPL (Figure 4D).

**Fig. 4.**
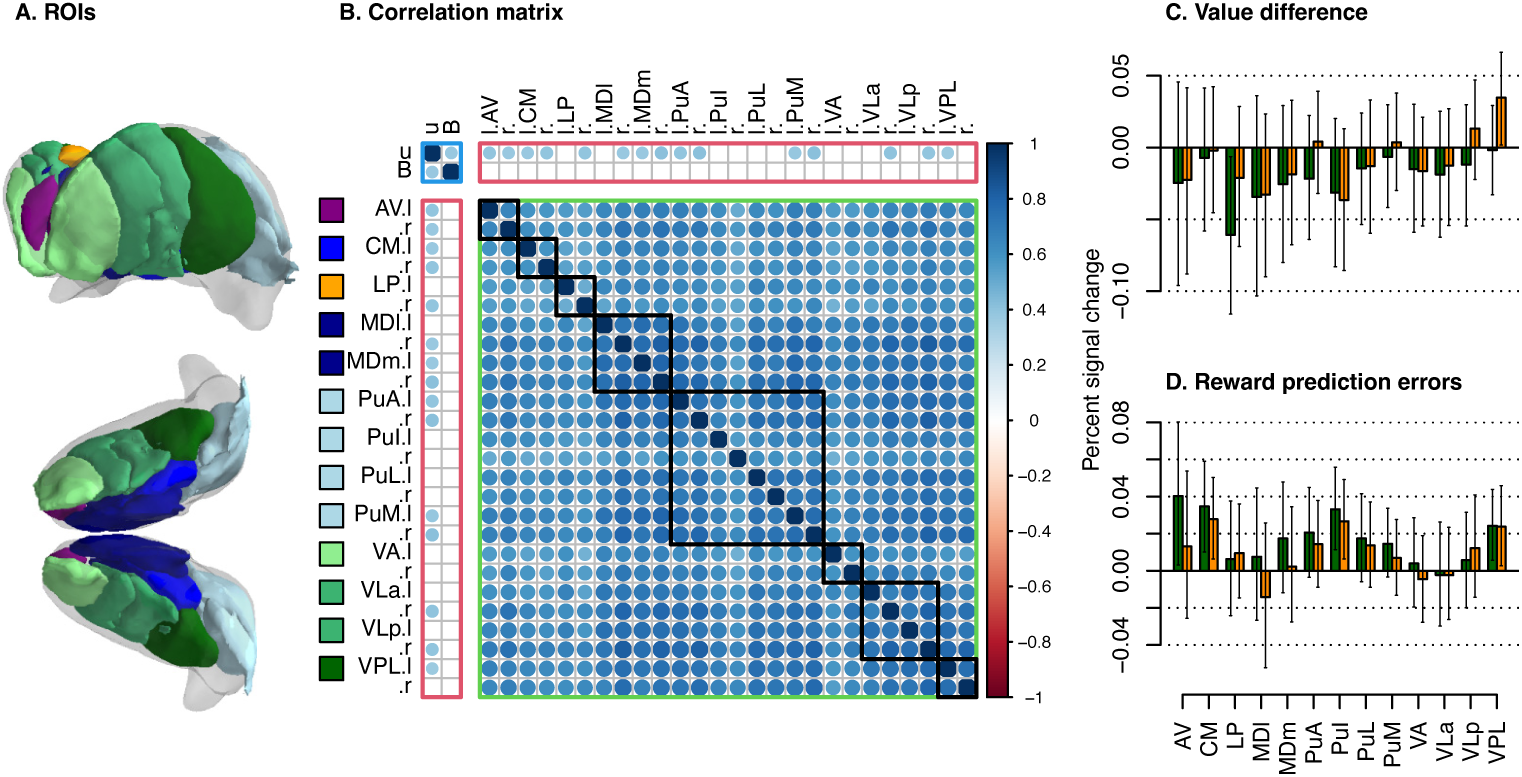
Joint model fit to the thalamus ROIs. A. Illustration of the ROIs, viewed from the front-left (top) and bottom (bottom). Meshes were generated by first warping all individual-level delineations to MNI-space, and subsequently running the marching cube algorithm on the across-participant mean in MNI-space. For comparison, the MASSP delineation of the thalamus is illustrated in transparent white. B. Group-level correlation matrix, which is split into behaviour-behaviour relations (outlined by a blue rectangle), brain-brain relations (red), and brain-behaviour relations (green). Subregions belonging to the same nuclei are clustered along the diagonal with black squares. Only credible corre-lations are shown; non-credible correlations are displayed as empty squares. Relations are considered credible when the 95% credible interval of the correlation coefficient does not cover 0. All parameters are related to the speed-accuracy trade-off contrast: Its effect on urgency (u), threshold (B), and the BOLD contrast in the ROIs. C and D. Group-level estimates of within-participant brain-behaviour relations of value learning and reward prediction errors. Barplots show the percentage signal change per unit change in value difference (C) and reward prediction errors (D), for each region of interest. Green and orange bars depict the left and right hemisphere, respectively. Error bars indicate 95% credible intervals.

In a third and final joint model, we zoomed in on the striatum. Unlike the thalamus, the human striatum is a relatively homogeneous structure, without clear internal cytoarchitectural or immunohistochemical boundaries between the dorsal and ventral striatum (e.g., Haber & Knutson, 2010). However, it has long been argued to be functionally specialised in multiple zones (e.g., Pauli, O’Reilly, Yarkoni, & Wager, 2016), with distinct afferent projections (del Rey & García-Cabezas, 2023; Haber & Knutson, 2010). Here, we used the recently developed second iteration of MASSP (Bazin et al., under review) to delineate the striatum into three separate parts: the nucleus accumbens (nAcc), putamen (Pu^1^), and caudate (Cau) (Figure 5A). We would like to point out that the nAcc in MASSP was delineated using a perpendicular line at the base of the internal capsule, which may result in the inclusion of an area that is not fully restricted to the nAcc. The joint model fit to the timeseries of these subregions is shown in Figure 5B-D (see Table A3 for numerical estimates). The brain-behaviour association relating to speed-accuracy trade-off settings was strongest in the dorsal striatum (Pu and Cau), and only credible in the right (but not left) nAcc. As expected, reward prediction error processing was clearest in the nAcc, but also detectable in both the Pu and the Cau (Figure 5C). A positive association between the size of the BOLD responses and the size of value differences was found in the Pu, and interestingly, a *negative* association in the Cau (and no association in the nAcc) (Figure 5D).

**Fig. 5.**
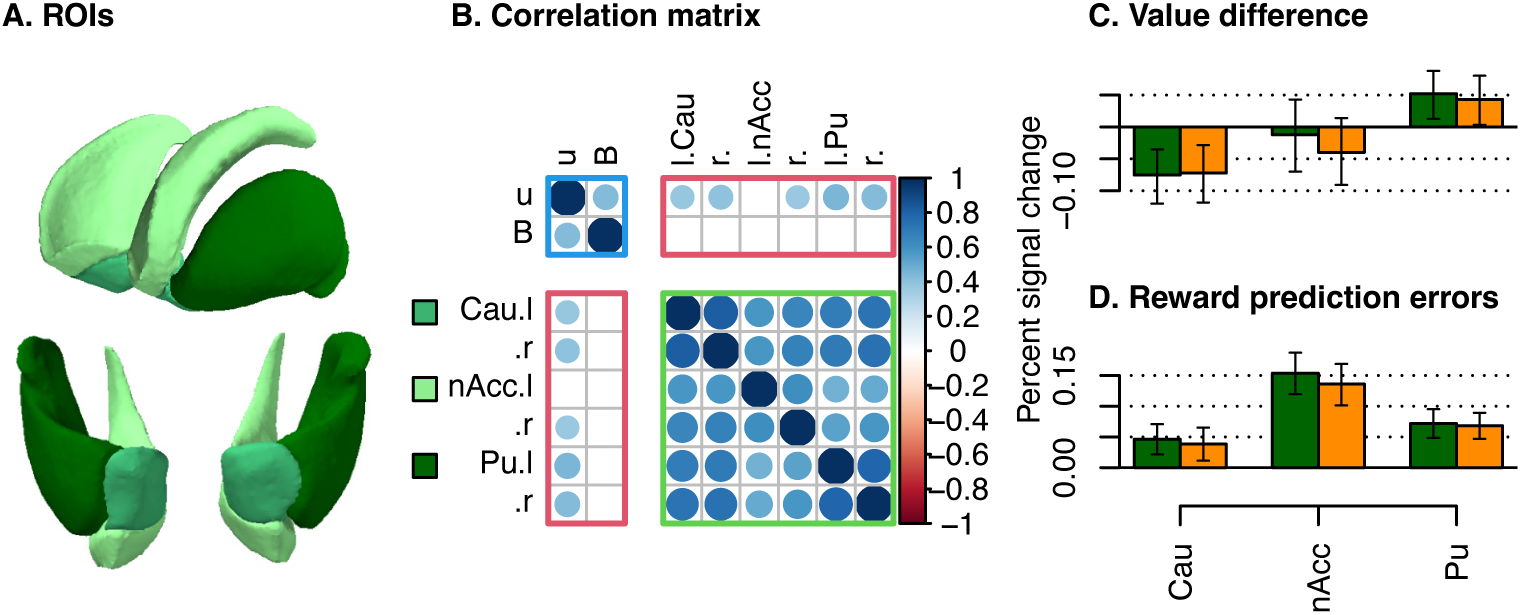
Joint model fit to the striatum ROIs. A. Illustration of the ROIs, viewed from the front-left (top) and bottom (bottom). B. Group-level correlation matrix, which is split into behaviour-behaviour relations (outlined by a blue rectangle), brain-brain relations (red), and brain-behaviour relations (green). Only credible correlations are shown; non-credible correlations are displayed as empty squares. Relations are considered credible when the 95% credible interval of the correlation coefficient does not cover 0. All parameters are related to the speed-accuracy trade-off contrast: Its effect on urgency (u), threshold (B), and the BOLD contrast in the ROIs. C and D. Group-level estimates of within-participant brain-behaviour relations of value learning and reward prediction errors. Barplots show the percentage signal change per unit change in value difference (C) and reward prediction errors (D), for each region of interest. Green and orange bars depict the left and right hemisphere, respectively. Error bars indicate 95% credible intervals.

## 3 Discussion

In this study, we use joint models to understand the brain-behaviour relations between subcortical regions and decision-making and learning. With tailored methods, includ-ing ultra-high field 7 T fMRI, decision making and instrumental learning were jointly studied in a single paradigm and corresponding cognitive model, in a Bayesian hierarchical joint modelling framework in which brain-behaviour relationships were reciprocally informed by all modalities of data. The resulting joint models revealed that the Str (and particularly the dorsal Str) was involved in choice strategy settings; however, contrary to previous reports, they demonstrated a relation with urgency, rather than response caution. Next, they revealed value-related processing, but not reward prediction error processing, in the substantia nigra. Finally, within the Str, value-related processing was demonstrated to show BOLD responses with opposite polarities in the caudate and putamen.

Our results indicate that subcortical regions may contribute to strategic control of choice behaviour through urgency, rather than response caution settings, which has been argued previously (Forstmann et al., 2008; Van Maanen et al., 2011). In part, this may arise from the use of the RL-ARD, which is able to dissociate urgency from response caution adjustments, which themselves correlate (e.g., Figure **??**). The implication of urgency adjustments corroborates earlier studies based on neural recordings the basal ganglia in monkeys (Thura, Cabana, Feghaly, & Cisek, 2022; Thura & Cisek, 2017), as well as fMRI evidence using an expanded judgment task (Van Maanen, Fontanesi, Hawkins, & Forstmann, 2016). The dominance of the right hemisphere in these relations is consistent with previous studies (Forstmann et al., 2008; Van Maanen et al., 2011, 2016), and may be related to the right-lateralised response inhibition networks (Cieslik, Mueller, Eickhoff, Langner, & Eickhoff, 2015; Hung, Gaillard, Yarmak, & Arsalidou, 2018; Isherwood, Keuken, Bazin, & Forstmann, 2021).

While our model-based approach is able to dissociate between urgency and threshold, the concept of urgency itself is not singular, as multiple cognitive processes may contribute to or correlate with urgency signals. For example, the CM plays an important role in modulating arousal (Motelow & Blumenfeld, 2014), which is related to urgency (Murphy, Boonstra, & Nieuwenhuis, 2016) (and covaries with reward prediction errors; Anderson et al., 2003; Small et al., 2003). The MD has been implicated in various types of memory processing, including object-reward association memory (e.g. Aggleton & Mishkin, 1983a, 1983b; Gaffan & Parker, 2000; Li, Inoue, Nakagawa, & Koyama, 2004; Zola-Morgan & Squire, 1985). The role of the MD may be to prepare the memory processes required for the subsequent value-based decision, and such preparations could start earlier under speed stress. Some evidence also suggests a role for the AV in modulating cortical plasticity and memory formation (Child & Benarroch, 2013). The involvement of the RN and VTA in urgency has, to our knowledge, not been demonstrated before, but may be related to earlier studies that demonstrate these regions’ involvement in conflict resolution, which is potentially caused by the conflicting instructions of the speed and accuracy requirements (Isherwood et al., 2023, 2024). The discovery of brain regions that correlated with urgency settings, and their function, can help us theorise about potential confounds of urgency that are difficult to derive based on behavioural studies alone. Model-based analyses should be combined with clever experimental design and manipulations to disentangle the influences of various confounding factors to estimated brain-behaviour relations.

Our results further indicated value-related processing in the Str, but with opposite polarities in the Cau compared to the Pu. This striking result might reflect a gradient of functional specialisation related to value differences. Alternatively, recent research has shown that neural activity in the dorsal Str can elicit vasoconstriction and *negative* BOLD responses, implying that our finding of negative BOLD responses could nonetheless indicate increased neural activity (Cerri et al., 2024). Note that value differences, in the present design, are confounded by other factors, which importantly includes difficulty: A choice based on two stimuli which differ in their value is easier compared to stimuli with similar values. Additional confounding factors include salience and arousal effects (see also O’Doherty, 2014). Disentangling the influence of these factors requires specific experimental designs in future studies.

We further found various subcortical regions in which BOLD responses covaried with reward prediction error sizes. While amygdalar and striatal involvement in reward prediction error coding are well-documented (e.g. Anderson et al., 2003; Small et al., 2003), the Cl and GPe received less attention in the literature. Recently, electrophysiological recordings in rodents identified a neural subpopulation encoding reward prediction errors in the GPe (Farries, Faust, Mohebi, & Berke, 2023). To some extent, these may arise also from covariates, such as perceived saliency (Kutlu et al., 2021, 2022). The Cl involvement might indicate a functional role similar to the Amg in terms of arousal and salience detection (Madden et al., 2022). It is becoming increasingly clear that dopamine signals can be detected in a wider range of behaviours than classical reward prediction errors, and can also signal sensory and motor features (for review, Gershman et al., 2024). Under the generalised prediction error framework (Gardner, Schoenbaum, & Gershman, 2018), they are argued to also indicate errors in the sensory world model, and are used to improve the world model. Consequently, a wide set of brain regions are likely involved in the processing of these predictions errors.

Contrary to some previous reports, we did not find evidence for dopaminergic midbrain involvement in reward prediction error encoding. A long history of animal recordings has implicated especially the VTA in reward prediction error processing (e.g. Montague et al., 1996; Schultz, 1986; Schultz et al., 1993, 1997; Watabe-Uchida et al., 2017), which has partially been supported in humans using fMRI (D’Ardenne, McClure, Nystrom, & Cohen, 2008; Fontanesi, Gluth, Rieskamp, & Forstmann, 2019; Hauser, Eldar, & Dolan, 2017; Pauli et al., 2015; Zhang, Larcher, Misic, & Dagher, 2017), but not consistently (see Garrison, Erdeniz, & Done, 2013, for a meta-analysis). A variety of factors has been argued to contribute to this discrepancy, including variability in the anatomical masks (Fontanesi, Gluth, Rieskamp, & Forstmann, 2019) and limited statistical power, as detailed in the introduction section. On the contrary, in the present study, the joint models were sufficiently powerful to identify value-related processing in the SN. Perhaps the discrepancy between the electrophysiology and BOLD findings is the result of a much more fundamental difference in methodology: While electrophysiology suggests reward prediction errors in the dopaminergic midbrain are encoded in spiking activity, BOLD responses have long been argued to correlate more strongly with synaptic activity (Goense & Logothetis, 2008; Hall, Howarth, Kurth-Nelson, & Mishra, 2016; Logothetis, Pauls, Augath, Trinath, & Oeltermann, 2001), which could indicate local processing as well as input to a region. It has often been argued, for example, that the striatal BOLD responses are a result of dopamine release caused by dopaminergic midbrain neural spiking (Ferenczi et al., 2016; Lohrenz, Kishida, & Montague, 2016). Intriguingly, since reward prediction errors are defined as the difference between obtained and expected reward, a region that calculates prediction errors needs expected reward (or value) as an input. This may explain why the SN BOLD responses were sensitive to value processing, but not reward prediction errors.

In conclusion, this study revealed various human subcortical underpinnings of decision making and learning. It uncovered new brain-behaviour relations (e.g., thalamic nuclei in urgency settings, GPe in reward prediction error processing), and refined previous work (e.g., functionally specialised zones along the anterior-posterior axis of the Str in value processing). It also demonstrates feasibility and value of the combination of joint modelling and tailored fMRI methods in progressing our understanding of the human subcortex in cognitive processes.

## 4 Methods

### 4.1 Participants

Thirty-seven healthy volunteers (mean age 27 years old [SD 6 years, range 19–39 years old], 20 females) were recruited via local advertisement. The study was approved by the Ethics Review Board of the Faculty of Social and Behavioral Sciences of the University of Amsterdam (reference: 2021-BC-13146) and the Regional Committees for Medical and Health Related Research Ethics of Central Norway (reference: 116630). All participants gave written informed consent prior to the onset of the study. All participants were screened for MRI safety, had normal or corrected-to-normal vision, and no history of psychiatric and neurological illness. All participants participated in five scanning sessions as part of a larger project; here, we report and analyse two of these sessions.

### 4.2 Paradigm

The experimental paradigm made use of an instrumental learning task (Frank, Seeberger, & O’Reilly, 2004) with a cue-based speed–accuracy trade-off manipulation (Miletić et al., 2021, see Figure 1). In every trial of the task, the participants made a decision between two abstract symbols, each associated with a fixed reward probability that is unknown to the participants. One choice option always had a higher probability of being rewarded than the alternative option. The participants received feedback in the form of points after each choice, which the participants can use to learn which symbols have the highest reward probabilities.

Prior to each trial, the participants were presented with a cue to instruct them to emphasise either response speed (‘SPD’) or accuracy (‘ACC’) on the upcoming trial. Speed and accuracy cues were randomly interleaved. On speed trials, participants had to respond within 700 ms to be eligible for a reward; on accuracy trials, participants had to respond within 1.5 s. After each choice, participants received two types of feedback: Firstly, the outcome of the choice (+0 or +100 points), and secondly, the actually obtained reward. If the participants responded in time (1.5 s in ‘ACC’ trials, 0.7 s in ‘SPD’ trials), their reward was equal to the outcome of the choice. If they responded too late, the participants were penalised with −100 points, irrespective of the outcome of the choice. The presentation of both the outcome of the choice and the actual reward allowed participants to both learn from the outcome of their choice as well as from their response timing.

In total, participants performed 342 trials divided over three runs. Each trial took 8.28 s (corresponding to 6 volumes; see below). Each run consisted of 3 new stimuli sets, that differed in their reward probabilities (80%/20%, 70%/30%, 60%/40%, respectively, for the three stimuli sets within each block) and therefore difficulty. Event timing was jittered to decorrelate the BOLD response design matrix, by pseudo-randomly sampling the duration of each fixation cross from 0.5 s, 1 s, 1.5 s and 2 s. Additionally, 10% null trials were included, during which the screen remained empty for 8.28 s.

### 4.3 Cognitive model specification

The behavioural data were modelled with the reinforcement learning advantage racing diffusion (RL-ARD) model (Miletić et al., 2021), which is an instance of the broader class of combined reinforcement learning evidence accumulation models (RL-EAMs; Miletić, Boag, & Forstmann, 2020). The RL-ARD conceptualises decision making as a race between accumulators, each accumulating evidence for one choice option. The first accumulator to reach a common threshold-level of evidence *a* triggers the motor processes that execute the decision. The time to respond equals the time to reach the threshold, plus an intercept *t*0 that corresponds to the time required for early perception encoding and response execution.

In the RL-ARD, each accumulator accumulates the *advantage* of one choice option over another option. Specifically, the rate of evidence accumulation (the drift rate *v*) of each accumulator depends on three terms: an evidence-independent base rate *u* (urgency); the advantage of one choice option of the other option, weighted by free parameter *w_d_*; and the total amount of evidence, weighted by free parameters *w_s_*. ‘Evidence’ in this model is based on Q-values, which represent the participant’s internal belief about how rewarding each choice option is. For two-choice tasks such as in the present study, the drift rates for the two accumulators are:

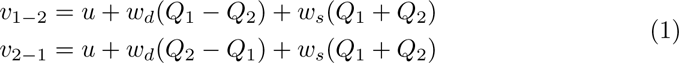

where *Q_i_* is the Q-value for choice alternative *i*, which are updated after every trial according to a simple delta rule:

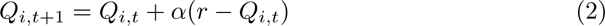

where *t* is the trial number, *r* is the obtained reward (in this specific experimental paradigm, the ‘outcome’), and *α* a free parameter known as the learning rate.

To model the effect of the SAT manipulation, we allowed both the *V*_0_ and *a* param-eters to vary freely between the speed and accuracy conditions, based on our earlier work (Miletić et al., 2021).

In total, the RL-ARD has eight free parameters: two evidence-independent base rate *u_acc_*and *u_spd_*, weights on the difference and sum of the evidence *w_d_* and *w_s_*, non-decision time *t*0, learning rate *α*, and two thresholds *a_spd_*and *a_acc_*. Instead of estimating a *u* parameter for each condition separately, we estimated the across-condition mean *u* and difference *u_spd−acc_* parameters (hence, *u_spd_* = *u* + *u_spd−acc_* and *u_acc_* = *u* − *u_spc−acc_*), and similarly, we estimated an across-condition mean *a* and difference *a_spd−acc_* parameter. The direct estimation of the between-condition differences in these parameters facilitates estimation of covariance with neural model parameters, which are detailed below.

### 4.4 MRI data acquisition

In multiple sessions, participants were scanned in a MAGNETOM ‘Terra’ 7 T MRI system (Siemens Healthineers, Germany) with a 32-channel phased array head coil (Nova Medical Inc, USA). The first session contained two anatomical scans: A multi-echo gradient recalled echo (GRE) and an MP2RAGE, both at 0.75 mm isotropic resolution. For the MP2RAGE, we used the following parameters: repetition time (TR) = 4.3 s, inversion times (TI_1,2_) = 840 ms and 2370 ms, flip angles (FA_1,2_) = 5*^◦^* and 6*^◦^*, echo time (TE) = 1.99 ms, field of view (FOV) = 240 × 240 × 168 mm, band-width = 250 Hz*/*px. For the GRE, the following parameters were used: TR = 31.0 ms, TE_1-4_ = 2.51, 7.22, 14.44 and 23.23 ms, FA = 12*^◦^*, FOV = 240 × 240 × 168 mm. In the remainder of this anatomical session, resting state data was collected that is not of interest for the current study.

The second session contained three functional runs with the task paradigm. A single echo echo planar imaging (EPI) sequence was used designed by the CMRR (https://www.cmrr.umn.edu/multiband/), with parameters based on our previous studies (De Hollander, 2017; Miletić, Bazin, et al., 2020) to tailor the sequence for the subcortex: 1.5 mm isotropic resolution, TE 14 ms, TR 1.38 s, partial Fourier 6*/*8, in-plane acceleration (GRAPPA) 3, multiband 2, bandwidth 1446 Hz*/*px, phase encoding direction A*>*P, FOV = 192 × 192 × 132 mm. In contrast to our previous work, we included a multiband factor of 2 in the protocol. Pilot testing indicated that on this MRI system, the increase in statistical power obtained through the increase in number of volumes (due to the lower TR with multiband acquisition) outweighed the loss in SNR (even in subcortical areas) for statistical testing purposes.

Each run consisted of 754 volumes (17 minutes 56 seconds). Immediately after each run, we collected 5 volumes of the same protocol with opposite phase encoding direction (P*>*A), which was used for susceptibility distortion correction purposes. Finally, at the end of the functional session, a low-resolution 1 mm MP2RAGE scan was acquired for co-registration purposes, using the same parameters as in the anatomical session.

During functional runs, physiological data on the participant’s heart rate and respiration were acquired using a photoplethysmograph (with sampling frequency 200 Hz) and respiratory belt (with sampling frequency 50 Hz), respectively. In six runs (two in one participant, one in another participant, and three in a third participant), recording of physiological data failed due to technical reasons.

### 4.5 Anatomical masks

We used the multi-contrast anatomical subcortical structure parcellation (MASSP) algorithm (Bazin et al., 2020) to obtain participant-specific anatomical masks of 17 subcortical structures. MASSP relies on multiple contrasts; here, we used quantified susceptibility (QSM) values, the longitudinal relaxation rates (R1), and effective transverse relaxation rates (R2^*^). R1 values were computed based on the MP2RAGE data using a look-up table (Marques et al., 2010). R2^*^ values were computed by least squares fitting of a mono-exponential decay function to the four echoes of the GRE data. QSM values were obtained using the phase maps of the last three echoes of the GRE data (Caan et al., 2019) with TGV-QSM (Langkammer et al., 2015). In both cases, LCPCA denoising (Bazin et al., 2019) was performed beforehand on the 8 images of the GRE (4 magnitude and 4 phase). Prior to estimating R2^*^ and QSM, the GRE data were brought into MP2RAGE-space by co-registration of the first GRE echo (magnitude image) to the second inversion of the MP2RAGE, using a rigid transformation in ANTs.

The MASSP algorithm combines shape, location, and R1, R2^*^, QSM value priors to delineate the following 17 subcortical structures in an individual’s data: Amygdala (Amg), claustrum (Cl), fornix (fx), the external and internal segments of the globus pallidus (GPe, GPi), internal capsule (ic), periaqueductal grey (PAG), pedunculopontine nucleus (PPN), red nucleus (RN), substantia nigra (SN), subthalamic nucleus (STN), striatum (Str), thalamus (Tha), ventral tegmental area (VTA), and the lateral, third, and fourth ventricles (LV, 3V, 4V). For all regions except fx, 3V and 4V, separate masks were obtained for both hemispheres. Here, we only focus on the gray matter structures, and thus excluded the internal capsule, fornix, and ventricles from the ROI analyses below; totalling 12 ROIs bilaterally.

Like in Miletić, Bazin, et al. (2022), we trained the MASSP algorithm on renormalised versions of the quantitative contrasts using a fuzzy C-means clustering of intensities, and linearly interpolating between cluster centroids. We also registered the data to the MASSP atlas in two successive steps. These alterations compared to the original MASSP implementation (Bazin et al., 2020) led to small parcellation improvements for some structures.

To segment the thalamus into individual nuclei, we used the thalamic segmentation tool *segmentThalamicNuclei.sh* as part of *freesurfer* 7.2.0. The segmentation applies a probabilistic atlas that was built using a combination of *in vivo* and *ex vivo* data (Iglesias et al., 2018). The segmentation is performed in subject space with the T1w contrast after running the freesurfer pipeline (recon-all) as part of *fmriprep* (see below). The tool outputs discrete segmentations at a resolution of 0.5 mm, which were resampled to 1.5 mm resolution with linear interpolation.

### 4.6 fMRI preprocessing

Results included in this manuscript come from preprocessing performed using *fMRIPrep* 20.2.0 (Esteban et al. (2018); Esteban et al. (2019); RRID:SCR 016216), which is based on *Nipype* 1.5.1 (Gorgolewski et al. (2011); Gorgolewski et al. (2018); RRID:SCR 002502).

#### 4.6.1 Anatomical data preprocessing

A total of 2 T1-weighted (T1w) images were found within the input BIDS dataset: One at 0.75 mm resolution from the anatomical session, and one at 1.0 mm resolution from the session including the task paradigm. Both were corrected for intensity non-uniformity (INU) with N4BiasFieldCorrection (Tustison et al., 2010), distributed with ANTs 2.3.3 (Avants, Epstein, Grossman, & Gee, 2008, RRID:SCR 004757). The T1w-reference was then skull-stripped with a *Nipype* implementation of the antsBrainExtraction.sh workflow (from ANTs), using OASIS30ANTs as target template. Brain tissue segmentation of cerebrospinal fluid (CSF), white-matter (WM) and gray-matter (GM) was performed on the brain-extracted T1w using fast (FSL 5.0.9, RRID:SCR 002823, Zhang, Brady, & Smith, 2001). A T1w-reference map was computed after registration of 2 T1w images (after INU-correction) using mri robust template (FreeSurfer 6.0.1, Reuter, Rosas, & Fischl, 2010). Brain surfaces were reconstructed using recon-all (FreeSurfer 6.0.1, RRID:SCR 001847, Dale, Fischl, & Sereno, 1999), and the brain mask estimated previously was refined with a custom variation of the method to reconcile ANTs-derived and FreeSurfer-derived segmentations of the cortical gray-matter of Mindboggle (RRID:SCR 002438, Klein et al., 2017). Volume-based spatial normalization to one standard space (MNI152NLin2009cAsym) was performed through nonlinear registra-tion with antsRegistration (ANTs 2.3.3), using brain-extracted versions of both T1w reference and the T1w template. The following template was selected for spatial normalization: *ICBM 152 Nonlinear Asymmetrical template version 2009c*[Fonov, Evans, McKinstry, Almli, and Collins (2009), RRID:SCR 008796; TemplateFlow ID: MNI152NLin2009cAsym].

#### 4.6.2 Functional data preprocessing

For each of the 3 BOLD runs found per subject, the following preprocessing was performed. First, a reference volume and its skull-stripped version were generated by aligning and averaging 1 single-band references (SBRefs). A B0-nonuniformity map (or *fieldmap*) was estimated based on two EPI references with opposing phase-encoding directions, with 3dQwarp Cox and Hyde (1997) (AFNI 20160207). Based on the estimated susceptibility distortion, a corrected EPI reference was calculated for a more accurate co-registration with the anatomical reference. The BOLD reference was then co-registered to the T1w reference using bbregister (FreeSurfer) which implements boundary-based registration (Greve & Fischl, 2009). Co-registration was configured with six degrees of freedom. Head-motion parameters with respect to the BOLD reference (transformation matrices, and six corresponding rotation and translation parameters) are estimated before any spatiotemporal filtering using mcflirt (FSL 5.0.9, Jenkinson, Bannister, Brady, & Smith, 2002). BOLD runs were slice-time cor-rected using 3dTshift from AFNI 20160207 (Cox & Hyde, 1997, RRID:SCR 005927). First, a reference volume and its skull-stripped version were generated using a custom methodology of *fMRIPrep*. The BOLD time-series (including slice-timing correction when applied) were resampled onto their original, native space by applying a single, composite transform to correct for head-motion and susceptibility distortions. These resampled BOLD time-series will be referred to as *preprocessed BOLD in original space*, or just *preprocessed BOLD*. Several confounding time-series were calculated based on the *preprocessed BOLD* : framewise displacement (FD), ‘DVARS’ (the spatial standard deviation of difference images), and three region-wise global signals. FD was computed using two formulations following Power (absolute sum of relative motions, Power et al. (2014)) and Jenkinson (relative root mean square displacement between affines, Jenkinson et al. (2002)). FD and DVARS are calculated for each functional run, both using their implementations in *Nipype* (following the definitions by Power et al., 2014). The three global signals are extracted within the CSF, the WM, and the whole-brain masks. Additionally, a set of physiological regressors were extracted to allow for component-based noise correction (*CompCor*, Behzadi, Restom, Liau, & Liu, 2007). Principal components are estimated after high-pass filtering the *preprocessed BOLD* time-series (using a discrete cosine filter with 128 s cut-off) for the two *CompCor* variants: temporal (tCompCor) and anatomical (aCompCor). tCompCor components are then calculated from the top 2% variable voxels within the brain mask. For aCompCor, three probabilistic masks (CSF, WM and combined CSF+WM) are generated in anatomical space. The implementation differs from that of Behzadi et al. in that instead of eroding the masks by 2 pixels on BOLD space, the aCompCor masks subtract a mask of pixels that likely contain a volume fraction of GM. This mask is obtained by dilating a GM mask extracted from the FreeSurfer’s *aseg* segmentation, and it ensures components are not extracted from voxels containing a minimal fraction of GM. Finally, these masks are resampled into BOLD space and binarised by thresholding at 0.99 (as in the original implementation). Components are also calculated separately within the WM and CSF masks. For each CompCor decomposition, the *k* components with the largest singular values are retained, such that the retained components’ time series are sufficient to explain 50 percent of variance across the nuisance mask (CSF, WM, combined, or temporal). The remaining components are dropped from consideration. The head-motion estimates, calculated in the correction step, were also placed within the corresponding confounds file. The confound time series derived from head motion estimates and global signals were expanded with the inclusion of temporal derivatives and quadratic terms for each (Satterthwaite et al., 2013). Frames that exceeded a threshold of 0.5 mm FD or 1.5 standardised DVARS were annotated as motion outliers. All resamplings can be performed with *a single interpolation step* by composing all the pertinent transformations (i.e. head-motion transform matrices, susceptibility distortion correction when available, and co-registrations to anatomical and output spaces). Gridded (volumetric) resamplings were performed using antsApplyTransforms (ANTs), configured with Lanczos interpolation to minimise the smoothing effects of other kernels (Lanczos, 1964). Non-gridded (surface) resamplings were performed using mri vol2surf (FreeSurfer).

Many internal operations of *fMRIPrep* use *Nilearn* 0.6.2 (Abraham et al., 2014, RRID:SCR 001362), mostly within the functional processing workflow. For more details of the pipeline, see the section corresponding to workflows in *fMRIPrep*’s documentation.

### 4.7 Neural model specification: Whole-brain generalised linear models (GLMs)

The timeseries of the neural data were modelled using GLMs. In these GLMs, we modelled each voxel’s timeseries *y* as:

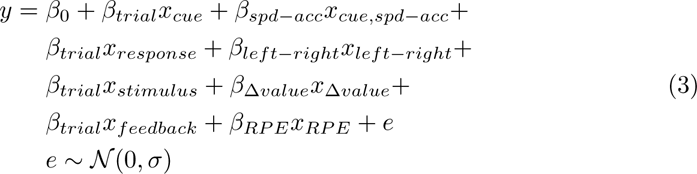

where every *β* is a parameter to be estimated, *x* are the timeseries of the experiment events convolved with the canonical double-gamma haemodynamic response function (HRF; Glover, 1999), and *σ* the residual variance. Note that we estimated a single *β_trial_* parameter to account for the shared effects of the presentations of cues, stimuli, and feedback, as well as the effects of motor responses (e.g., the effects of visual processing and overall motor preparation). In experimental paradigm, the effects of these event types cannot be disentangled from one another due to their rapid succession within a trial. Note, however, that the contrasts of interest are orthogonal to these events and can be estimated well.

Mirroring the cognitive model, we estimated a between-cue difference for the BOLD responses relating to the cue. Specifically, the regressor *x_cue,spd−acc_* was also modelled on the onset of the cue but shows a negative deflection for ‘ACC’ cues and a positive deflection for ‘SPD’ cues. As such, the corresponding *β_cue,spd−acc_* reflects the difference in ‘SPD’ over ‘ACC’ cues. Similarly, the *β_left−right_* parameter reflects the BOLD-contrast resulting from left compared to right motor responses. The corresponding *x_left−right_* regressor was modelled on the onsets of the button presses.

The regressors *x_stimulus_*and *x*_Δ*value*_ relate to the stimulus and stimulus value differences, respectively. The amplitude of the stimulus value regressor varied parametrically across trials, with the trial-by-trial amplitude determined based on the difference in Q-values (internal value representations) as estimated by the RL-ARD model. Similarly, the regressors *x_feedback_*and *x_RP E_* relate to the effects of the feed-back and the reward prediction error, respectively, which were simulated obtained from the RL-ARD. Both the value difference and reward prediction error regressors were demeaned per run, to orthogonalise them with respect to the stimulus and feed-back regressors. We included the temporal derivatives of all task regressors (note that these are not shown in Equation 3).

As control analyses, we first fit the GLM using a traditional two-stage mass-univariate approach, where a GLM is fit per voxel. In this approach, we first fit the RL-ARD to the behavioural data, and extracted trial-by-trial regressors per subject by simulating them from the RL-ARD model. Specifically, the model was used to simulate the task paradigm for 100 times, each time with a different set of RL-ARD parameters (randomly sampled from the posterior distributions). On each trial of the simulation, the difference in values of the two stimuli was calculated, and the mean of the stimulus value differences at each trial across the 100 simulations was used to determine the regressor’s amplitude. These stimulus value differences were then demeaned per run. The trial-by-trial height of the parametrically varying reward prediction error regressor was determined based on the same simulation of the RL-ARD (except now using the reward prediction error instead of the value differences), and this regressor was also demeaned per run.

To model physiological noise, we included a set of 18 regressors obtained using RETROICOR (Glover, Li, & Ress, 2000): 3th order phase Fourier expansion of the heart rate signal, 4nd order phase expansion of the respiration signal, and a 2nd order phase Fourier expansion of the interaction between heart rate and respiration (Harvey et al., 2008). Two additional regressors were used to model heart rate variability (HRV; Chang, Cunningham, & Glover, 2009), and respiratory volume per time unit (RVT; Birn, Smith, Jones, & Bandettini, 2008; Harrison et al., 2021). These physiological regressors were estimated using the PhysIO toolbox (Kasper et al., 2017) implemented in the TAPAS software package (Frässle et al., 2021). For six runs (one in a single participant, two in another participant, and three in a third participant), collection of the physiological data failed due to technical reasons. For these runs, the first 20 aCompCor components (Behzadi et al., 2007) were instead included in the design matrix. Additionally, for all participants 7 motion-related regressors were included (translation and rotation in three dimensions, plus the framewise displacement), and a set of discrete cosines to model low-frequency drifts. To model residual physiological noise, we also included a regressor with the mean signal within CSF, estimated by *fMRIprep*. Finally, we included a nuisance regressor to model the effect of response times using the *RTDur* approach (Mumford et al., 2024). This regressor is generated by convolving a boxcar function (starting at the onset of each stimulus, with the response time on that trial as duration) with the same haemodynamic response function as was used for the task-related regressors.

Prior to fitting the whole-brain GLM, the data were minimally smoothed using SUSAN (Smith & Brady, 1997, kernel size FWHM = 1.5 mm). Run-level GLMs were estimated using FSL FEAT (Woolrich, Ripley, Brady, & Smith, 2001), and afterwards the three run-level GLMs per participant were combined with a fixed effects analysis. Group-level models were estimated using FSL FLAME1+2 (Woolrich, Behrens, Beckmann, Jenkinson, & Smith, 2004). For the speed - accuracy cue contrast, the design matrix included both an intercept and two model-based parametrically varying parameters: the between-condition differences in the threshold parameter (speed - accuracy) and in the urgency parameter, which were z-scored. All group-level statistical parametric maps (SPMs) were corrected for the false discovery rate with the Benjamini-Hochberg procedure (FDR; *q <* 0.05). SPMs of the whole-brain results can be found in the Supplementary Results.

### 4.8 Joint models

The main analysis used joint models. In joint models, the cognitive model (RL-ARD) and the neural model (GLM) are estimated simultaneously (Turner, Forstmann, et al., 2017; Turner, Forstmann, & Steyvers, 2019; Turner, Palestro, et al., 2019; Turner et al., 2016; Turner, Wang, & Merkle, 2017). Furthermore, the joint models we employ assumed that the cognitive and neural parameters are multivariate normally distributed across subjects: *θ, β* ∼ *MV N* (*δ,* Σ). This assumption allows for estimation group-level mean parameters *δ* as well as correlations between parameters through the variance-covariance matrix Σ, and thereby allows for estimating which cognitive processes correlate with BOLD responses in which regions of interest (ROIs).

The variance-covariance matrix of a multivariate normal grows quadratically with the number of cognitive and neural parameters estimated. Therefore, we applied multiple restrictions to the participant-level models to retain feasibility of parameter estimation (Stevenson, Innes, Gronau, et al., 2024). Specifically, we made a distinction between estimating parameters *jointly* (i.e., both the group-level mean and the correlations between parameters of neural and cognitive models across individual) or *non-jointly* in which the group-level mean was estimated, but the no correlations were estimated.

Of the cognitive model, we estimated all parameters jointly. Figure **??** focuses on only the parameters related to the SAT manipulation. Of the neural model, we estimated the *β_cue,spd−acc_*, *β*_Δ*value*_, and *β_RP E_* parameters of interest jointly, as well as the *β_CSF_*and *β_RT_*nuisance parameters. We estimated only these latter nuisance parameters jointly as we hypothesised these could most strongly correlate with parameters of interest. All other neural parameters (including the temporal derivatives and the standard deviation of the errors) were estimated non-jointly.

Joint models were fit to neural data from the ROIs defined by MASSP and by the thalamus atlas. To obtain the signal per ROI, first, the mean timeseries within each ROI defined by MASSP was extracted from the unsmoothed functional data. The mean timeseries were rescaled to percent signal change, through division by the mean signal, multiplying by 100 and subtracting 100. To reduce the total number of parameters in the joint models, we first filtered the timeseries and design matrix by least square regression of the same set of confounds as used in the whole-brain GLMs (except for the CSF and RT regressors, which were estimated in the joint model), to reduce physiological noise and remove low-frequency drifts from the signal.

### 4.9 Bayesian estimation

To allow for estimation of whole-brain general linear models (GLM) of the neural data, we first fit the cognitive model to the behavioural data only. All model estimations were performed using a Bayesian particle Metropolis-within-Gibbs (PMwG) sampler (Gunawan, Hawkins, Tran, Kohn, & Brown, 2020; Stevenson, Innes, Boag, et al., 2024). The PMwG sampler strictly adheres to a hierarchical model in which group-level parameters and participant-level parameters are estimated simultaneously. The group level is modelled with a multivariate Gaussian distribution, which is updated using Gibbs sampling. At the participant level, chains are updated using a combination of particle sampling and Metropolis-Hastings. We followed earlier work (Stevenson, Innes, Boag, et al., 2024) by using four sampling stages. The first, pre-burn stage, was used to approximate the participant-level likelihood landscape for proposal distributions. The burn stage was run until the mean Gelman’s diagnostic (Gelman & Rubin, 1992) was below 1.1. The adaptation stage was used to collect samples to generate a distribution that allows for efficient proposal samples in the last stage, the sampling stage. This sampling stage was run until convergence (assessed using Gelman’s diagnostics and visual inspection of the chains). Samplers were run with three chains.

The priors on the group-level mean were Gaussian distributions. The mean and standard deviation of these priors of the cognitive model parameters were based on the posterior distributions described in (Miletić et al., 2021), which used the same task and model (experiment 2). The prior for *u* was set to *N* (2.5, 1), *B* to *N* (1.5, 1), *t*_0_ to *N* (0.15, 1), *w_d_* to *N* (2.25, 1), *w_s_* to *N* (0.5, 1), and *α* to *N* (0.12, 1). The *t*0, *w_s_*, and *w_d_* parameters were estimated on the log scale, and *α* on the probit scale. The prior for the contrasts of interest, *B_spd−acc_* and *u_spd−acc_*, were set to *N* (−0.5, 1) and *N* (0.5, 1), respectively (note that threshold and urgency should have opposite signs to allow for faster responding under speed stress: thresholds should *decrease*, but urgency should *increase*). Visual comparisons confirmed that the posteriors were not strongly influenced by the priors for the parameters of interest.

For the group-level (co-)variance matrix we used an inverse-Gamma — inverse-Wishart mixture prior with 2 degrees of freedom and a scale parameter of 0.3. These settings give rise to uniform priors on the correlations (Huang & Wand, 2013), for parts of the group-level covariance matrix that were allowed to covary.

To visualise the quality of model fit, we randomly sampled 100 parameter sets from the posterior distributions, and used these to simulate the experimental design. These posterior predictive distributions were then used to calculate the credible intervals by taking the range between the 2.5% and 97.5% quantile of the averages across participants for each behavioural measure (RT quantiles and accuracy).

Next, we fit the joint models, in which we used the same priors for the cognitive models, except we decreased the variance of the group-level means to 0.7 for *B_spd−acc_* and *u_spd−acc_*, and to 0.5 for the other parameters. The priors for the neural parameters were set to *N* (0, 0.1), except for the RT nuisance parameter, which was set to *N* (0, 0.001). Note that the amplitude of the RT nuisance regressor is much larger than the amplitudes of the other neural regressors, due to its duration being modelled (as opposed to using a stick function of 0.001 s). This also entails that the absolute parameter estimates are much smaller, hence, we also used a smaller variance for this parameter to stabilise estimation.

## Supplementary information

Supplementary results can be found in Appendix A.

## Acknowledgments

We thank Pål Erik Goa for supporting this study by facilitating data acquisition. We thank Sarah Habli, Lisbeth Røe, and Daniel R. Sokol-owski for their help collecting the data.

## Declarations

- Funding: SM, NS, SJSI, ACT, BUF were supported by an ERC-Consolidator grant (nr. 864750; BUF). AA was supported by the JPND (IronSleep; nr. 10510062110003) and the Dutch Organisation for knowledge and innovation in health, healthcare and well-being (ZonMw; nr. 09120012110015).
- Competing interests: P-LB is the owner of Full Brain Picture Analytics. The other authors do not have any competing interests to declare.
- Ethics approval and consent to participate: This study was approved by the Ethics Review Board of the Faculty of Social and Behavioural Sciences of the University of Amsterdam (reference: 2021-BC-13146) and the Regional Committees for Medical and Health Related Research Ethics of Central Norway (reference: 116630). Participants provided written informed consent prior to data collection.
- Data availability: Neural timeseries and behavioural data are available at https://osf.io/pc5bm/
- Code availability: All analysis code is available at https://osf.io/pc5bm/

**Fig. A1.**
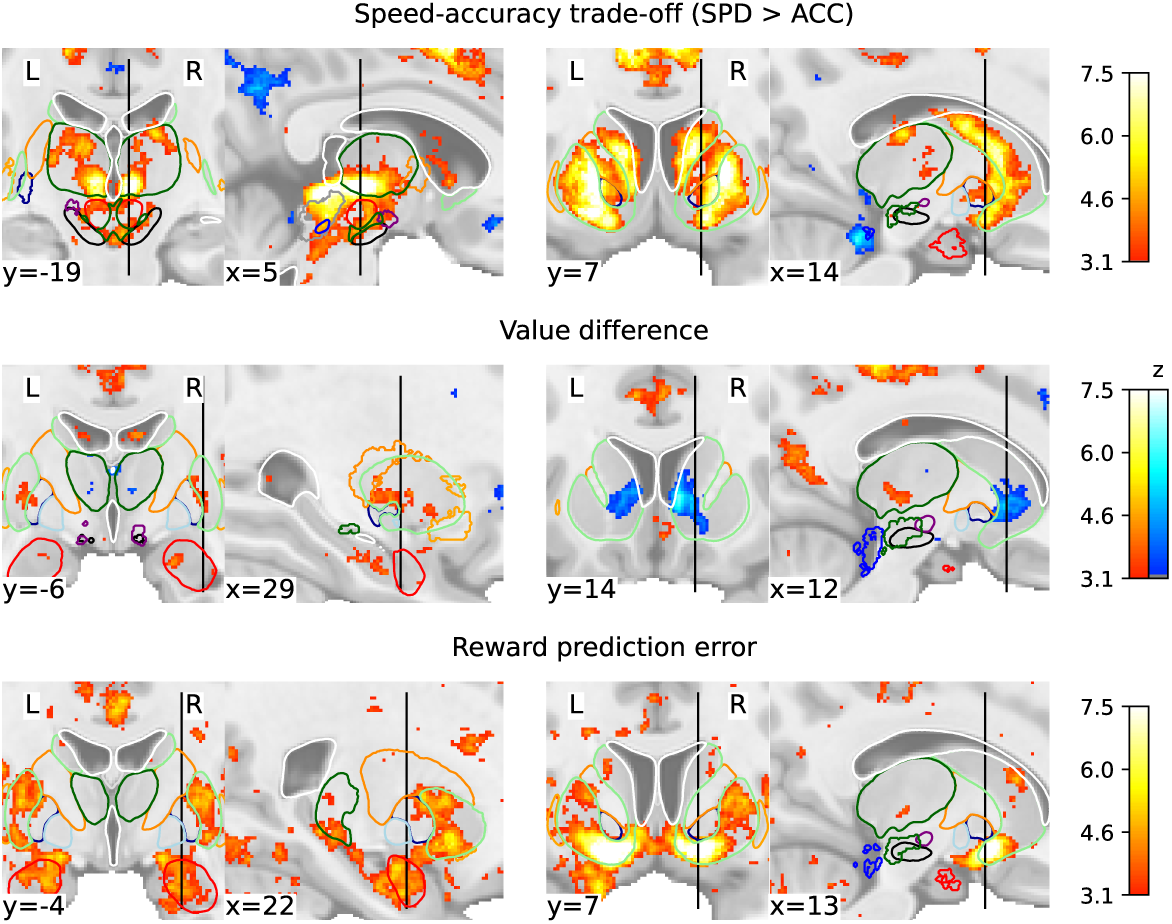
Subcortical statistical parametric maps. Top row focuses on the thalamic (left) and striatal (right) clusters in the speed >accuracy contrast. Middle row focuses on the striatal clusters in the value difference contrast. Bottom row focuses on the amygdalar (left) and ventral striatal (right) clusters in the reward prediction error contrast. Coordinates are in MNI152NLin2009cAsym (1 mm) space. Thresholds were determined using the FDR method (*q <* 0.05), with a minimum of *z* = 3.1 to prevent overly liberal thresholds. Contours illustrate the following regions: Amg (red), Cl (orange), fx (light gray), GPe (dark blue), GPi (light blue), ic (dark orange), PAG (grey), PPN (blue), RN (red), SN (black), STN (purple), Str (light green), Tha (dark green), lateral, 3rd and 4th ventricles (white), and VTA (dark green). Masks in MNI152NLin2009cAsym space were obtained by first applying the MASSP algorithm to each participant between age 18–40 of the data of Miletić, Bazin, et al. (2022). The resulting voxelwise probabilistic masks in subject space were then non-linearly warped to MNI space, after which the across-subject best label was determined per voxel. Contours delineate best labels.

## Appendix A Supplementary Results

### A.1 Whole-brain GLMs

#### A.1.1 Contrast 1: Speed–accuracy trade-off

First, we tested for regions that showed differential BOLD responses in the ‘SPD‘ and ‘ACC‘ conditions. In line with Forstmann et al. (2008), we found that the anterior parts of the dorsal striatum and preSMA showed larger BOLD responses to ‘SPD‘ compared to ‘ACC‘ cues. With the increased spatial resolution and minimal smoothing applied in the current study, we spatially located the BOLD responses specifically in the posterior part of the head of the caudate, as well as the anterior part of the putamen (see Figure A1 for more detail on the clusters in subcortex). In addition, we found BOLD clusters in the thalamus. Based on the atlas of thalamic subnuclei from Iglesias et al. (2018), the thalamic clusters corresponded to the ventral lateral (VL) nucleus, (38.3% of the cluster size in the left hemisphere, and 27.4% in the right hemisphere) and the ventral posteriolateral (VPL) nucleus (15.2% and 8.2% for the left and right hemisphere, respectively). Furthermore, 9-13% of the cluster size corresponded to the CM nucleus, and 16-18% to the MD nucleus.

#### A.1.2 Contrast 2: Value and difficulty

The second contrast was the value difference contrast. This contrast tests for regions in which the size of the BOLD responses parametrically covary with the size of the *difference* in expected value between the two choice options, as estimated using the cognitive model. Due to the nature of this task, however, the contrast of value differences conflates multiple cognitive factors (see also O’Doherty, 2014): The value difference determines the difficulty of the choice (higher value differences imply easier decisions), and thereby influences the evidence accumulation process (drift rates), and related, the (un)certainty regarding choices and the error rates. In part, the existing literature on perceptual decision making can help interpretation, as difficulty effects for cortical areas are well-documented (for a meta-analysis, see Keuken et al., 2014) and include the (anterior) insula, inferior frontal gyri, preSMA, and parietal regions that are more generally implicated in evidence accumulation such as the lateral intraparietal sulcus. Indeed, this network of regions is found in the value difference contrast.

Additionally, we found BOLD responses in various parts of the striatum: The putamen showed a positive correlation between value differences and BOLD responses, while the head of the caudate shows *negative* correlations.

#### A.1.3 Contrast 3: Reward prediction errors

The final contrast tested for regions showing a BOLD response of which the size covaried with the size of the RPEs, as estimated by the RL-ARD model. In line with the literature (for early studies, see e.g. McClure, Berns, & Montague, 2003; O’Doherty, Dayan, Friston, Critchley, & Dolan, 2003; O’Doherty et al., 2004), we found a clear BOLD cluster in ventral striatum (see Figure A1). The significant BOLD clusters covered many other regions across the entire brain, including the occipital lobe. BOLD responses that covary with RPEs in the occipital lobe have been found before in studies with comparable task paradigms (e.g., Hauser et al., 2017; Lefebvre, Lebreton, Meyniel, Bourgeois-Gironde, & Palminteri, 2017; Wimmer, Daw, & Shohamy, 2012, see also the corresponding statistical parametric maps on neurovault: collec-tions at https://identifiers.org/neurovault.collection:2007, neurovault.collection:2195, and neurovault.collection:2222, respectively). Furthermore, we found BOLD clusters in the ventral orbitofrontal cortex, another region often associated with value processing (Kringelbach & Rolls, 2004). Subcortically, we found clusters in the putamen, amygdala, and hippocampus.

**Fig. A2.**
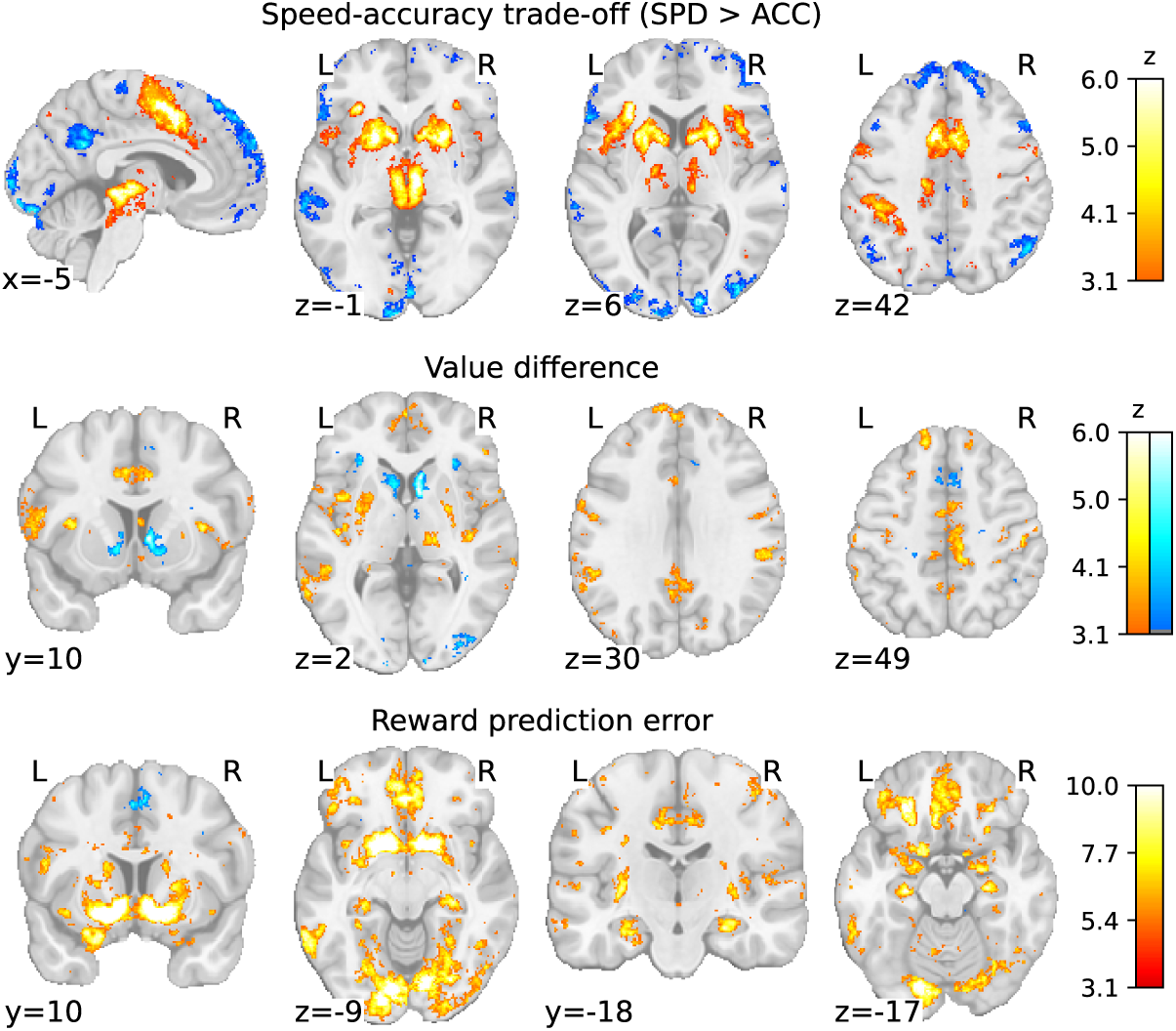
Results of the voxelwise whole-brain GLMs. The three rows show the three main contrasts of interest: The speed-accuracy trade-off (top), value differences (middle), and reward prediction errors (bottom).

**Table A1:**
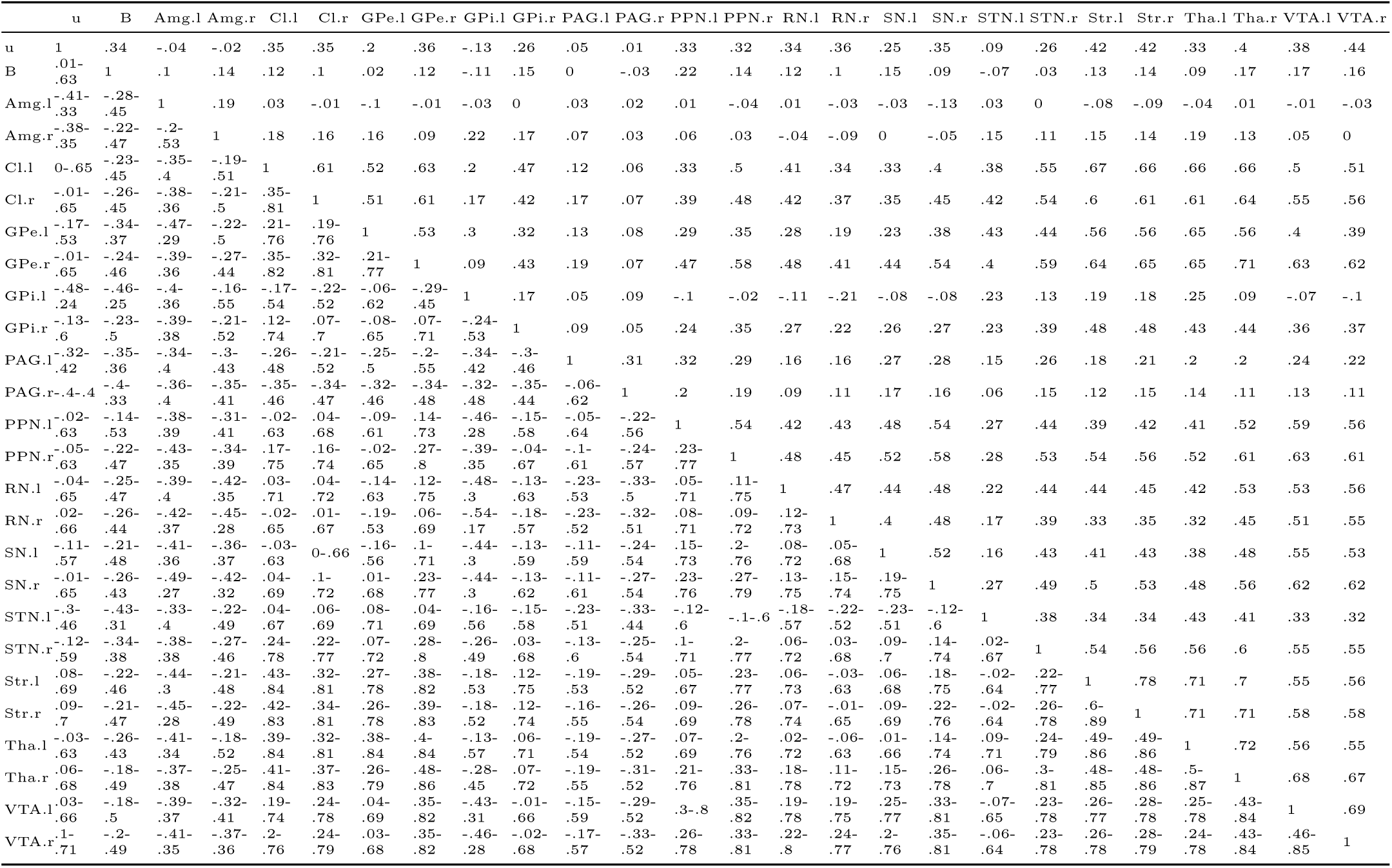
Correlation coefficients of parameters related to the speed-accuracy trade-off manipulation. Regions are from the MASSP delineations. Upper triangle displays means, lower triangle credible intervals.

**Table A2:**
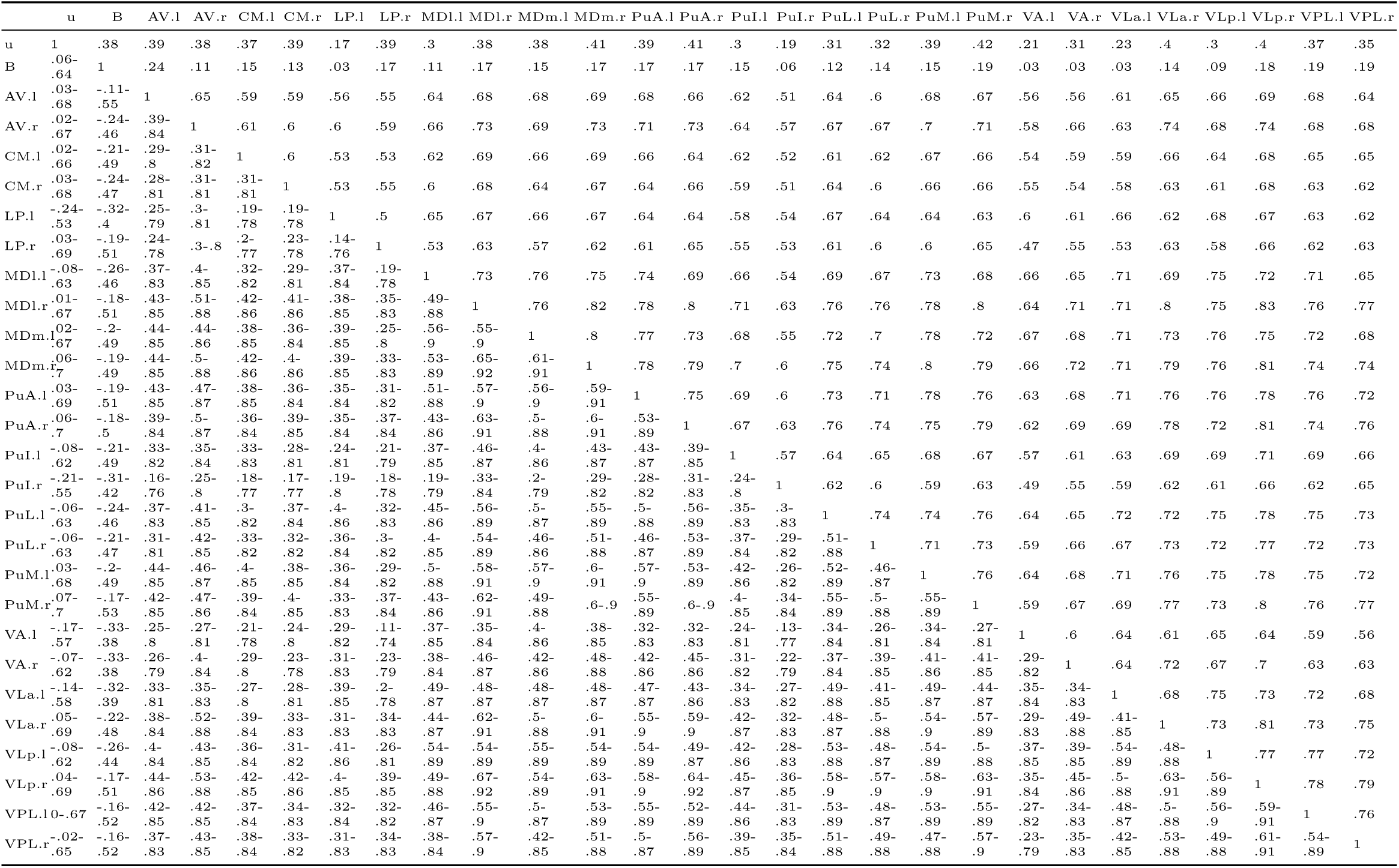
Correlation coefficients of parameters related to the speed-accuracy trade-off manipulation. Regions are from the thalamic nuclei delineations. Upper triangle displays means, lower triangle credible intervals.

**Table A3:** Correlation coefficients of parameters related to the speed-accuracy trade-off manipulation. Regions are subregions of the striatum. Upper triangle displays means, lower triangle credible intervals.

|  | u | B | Cau.l | Cau.r | nAcc.l | nAcc.r | Pu.l | Pu.r |
| --- | --- | --- | --- | --- | --- | --- | --- | --- |
| u | 1 | .44 | .38 | .4 | .23 | .37 | .45 | .44 |
| B | .11-.69 | 1 | .08 | .08 | .08 | .17 | .18 | .19 |
| Cau.l | .04-.67 | -.27-.43 | 1 | .83 | .59 | .66 | .7 | .73 |
| Cau.r | .07-.68 | -.27-.41 | .68-.92 | 1 | .58 | .68 | .71 | .74 |
| nAcc.l | -.17-.56 | -.28-.43 | .28-.81 | .28-.8 | 1 | .62 | .47 | .5 |
| nAcc.r | .04-.66 | -.19-.5 | .4-.83 | .44-.85 | .34-.82 | 1 | .54 | .58 |
| Pu.l | .11-.72 | -.18-.52 | .47-.86 | .48-.85 | .13-.73 | .23-.76 | 1 | .8 |
| Pu.r | .1-.71 | -.16-.51 | .53-.87 | .55-.88 | .17-.75 | .29-.79 | .64-.91 | 1 |

## Footnotes

1 Note that both the thalamic atlas and the second iteration of MASSP include ‘Pu’ as an abbreviation; the former referring to the Pulvinar, the latter to the Putamen. In this manuscript, Pu refers to the Putamen, and PuA, PuI, PuL and PuM to the various Pulvinar regions.

